# Spatial connectivity and marine disease dispersal: missing links in aquaculture carrying capacity debates

**DOI:** 10.1101/2023.06.20.545704

**Authors:** L. Schmittmann, K. Busch, L. C. Kluger

## Abstract

One major societal challenge is meeting the constantly increasing demand for (sea)food in a sustainable way. With marine aquaculture on the rise, it is crucial to define limits to aquaculture growth in order to ensure ocean health. Along these lines, the concept of aquaculture carrying capacity (CC) is increasingly intersected with the principles of the ecosystem approach to aquaculture. Its primary aims are to estimate sustainable production potential and limits of locally defined regions. However, the ocean is a fluid environment, subject to large- and small-scale dynamics, including ocean currents, tidal fluctuations, and human action. These dynamics introduce spatial connectivity between aquaculture sites and more distant ecosystems than considered in current CC estimates. We argue that far-reaching effects of aquaculture on the ocean, such as introduction and spread of invasive species and marine diseases, are thus underestimated when providing recommendations. Marine diseases can impact biodiversity, society, and overall ocean health and it is imperative to guide aquaculture development to reduce the risk of marine disease dispersal. We, therefore, suggest to embrace spatial ocean connectivity into the CC concept by using hydrodynamic modelling and dispersal simulations as high-throughput methods to estimate potential impact areas and provide risk assessments. In this work, we focus on the example of dispersing infectious diseases in bivalve farming and discuss ecological as well as social consequences of spatial connectivity. Both are applicable to a wide range of organisms and marine aquaculture systems internationally.

**Summary:** The concept of aquaculture carrying capacity (CC) aims at defining sustainable limits to aquaculture growth in order to ensure ocean health. Usually, estimations are based on locally defined regions and on the farm-scale. However, interactions of aquaculture with the ocean can have far-reaching effects, such as introduction and spread of invasive species and marine diseases. The ocean is a fluid environment, subject to large- and small-scale dynamics that introduce spatial connectivity between aquaculture sites and more distant ecosystems than considered in current CC estimates. We, therefore, suggest to embrace spatial ocean connectivity into the CC concept by using hydrodynamic modelling and dispersal simulations as high-throughput methods to estimate potential impact areas and provide risk assessments. Here, we focus on the example of dispersing infectious diseases in bivalve farming and discuss ecological as well as social consequences of spatial connectivity. Both are applicable to a wide range of organisms and marine aquaculture systems internationally.

## Definitions

We aim to examine the topic of aquaculture carrying capacity from an interdisciplinary perspective and use terms that can have differing meanings depending on context and discipline. Therefore, we provide definitions of key terms according to the use in this perspective article (marked in text in bold at first mention).

### Marine aquaculture

The breeding, rearing, and harvesting of marine plants and animals. Though this typically takes place in the ocean directly, the ocean-land interface is crossed when aquaculture input is received from specific facilities on land (e.g., hatcheries). Marine aquaculture is sometimes also referred to as mariculture.

### Ocean health^1^

Integrity, functionality, and resilience of the ocean ecosystem from a transdisciplinary perspective. Human societies are dependent on various ocean ecosystem services, and a sustainable use of, and interaction with the ocean must aim to secure ocean health.

### Aquaculture carrying capacity (CC)

Maximum level of aquaculture that a social-ecological system of a defined size can sustain before experiencing unacceptable changes to a state indicator.

### State indicator

State indicators are used to measure carrying capacity. They are variables to define maximum tolerable change induced by aquaculture before a system may experience undesirable impacts. Common state indicators are e.g., chlorophyll or oxygen concentration, or biomass of a non-cultured species.

### Dispersal (in the ocean)

The passive spreading of biotic or abiotic entities with water masses through ocean currents, or anthropogenic processes like ship transport, here termed “**anthropogenic dispersal**”.

### Spatial connectivity (in the ocean)

The connection and interaction between non-neighboring marine habitats or organisms through water movement. Water can move both naturally by ocean currents and anthropogenically, e.g., through ballast water transport.

### Marine diseases

In this context, we refer to infectious diseases of marine organisms caused by disease agents (such as bacteria, viruses, or protists) that can be transmitted directly between hosts, or via seawater.

### Wild and farmed organisms

Populations of organisms that occur naturally in the marine habitat, and populations of organisms that are intentionally cultured by humans for aquaculture purposes, respectively.

## Marine aquaculture and the carrying capacity concept

With declining wild fish and shellfish stocks and a simultaneously increasing demand for seafood, **marine aquaculture** is on the rise. The sector currently produces about 88 million tons of aquatic animals annually, which amounts to about 50% of the global production^2^. However, the aquaculture industry is criticized for its growth regardless of the consequences for **ocean health** (FAO 2022^2^ p. 18). The effects of aquaculture on surrounding ecosystems can take many forms, of which are depending on the farmed species, the environmental setting, and additional anthropogenic stressors^3–5^. The alteration of the physical-environmental space may have been most prominently discussed: be it through resource depletion^6^, changes in ocean currents^7,8^, release of waste products and nutrients^9^, shifts in the gene pool of wild organisms^10^, the introduction of potentially invasive species^11^, or erosion^12^.

All of these, and similar interactions of aquaculture with the ocean ecosystem may threaten biodiversity and ultimately what can be covered under the umbrella term “ocean health”: ecosystem integrity, functioning and resilience^1^. In turn, compromised ecosystem services affect human communities who are depending on the ocean for food supply or economy^1,13^. In overall, the aquaculture sector is challenged to find sustainable solutions as to not compromise ocean nor human health.

The concept of **aquaculture carrying capacity (CC)** has created much attention for debates on sustainable aquaculture growth. It aims at defining limits to production in order to create a “safe”^14^ operating space for aquaculture management^15,16^. The application of the ecological concept of CC to aquaculture settings emerged in the 1990s^17^. It is based on the assumption that an ecosystem can only sustain a certain level of biomass grown in culture, before unacceptable changes to the ecosystem occur, e.g., a decrease in biodiversity or physical space^15,16,18^.

Yet, what represents an unacceptable change to a system is easier defined theoretically, than put into practice. Different **state indicators** and the change thereof have been used to define thresholds for unacceptable change to a specific system as induced by aquaculture operations within that same setting^19^. Some approaches focus on physical production constraints of aquaculture: the location of farm sites and the availability of space (i.e. *physical* CC^20,21^), or the maximization of stocking density without compromising individual growth and oxygen concentrations (i.e. *production CC*^22^). Others discuss ecological concerns which acknowledge that marine aquaculture does not take place in an enclosed box but is embedded in a complex and dynamic ecosystem. Farms may, for example, cause aquaculture-external species to decline (i.e. *ecological CC*^23–26^), ecosystem integrity to be compromised^27^, or exceed the system’s capacity to deal with organic matter, nutrient and contaminant input (i.e., *assimilative CC*^14,28–30^). Fewer studies have looked at socio-economic considerations, such as social acceptance or willingness to pay, for guiding aquaculture expansion (i.e., *social CC*^31,32^). In the end, any limiting value for aquaculture expansion should necessarily be shaped by social debates, and processes to estimate CC should be multidimensional, iterative, inclusive and just^33^. Importantly, they should be backed on the governmental level^18,34,35^.

As such, the CC concept has found consideration in the ecosystem approach to aquaculture (EAA) that aims at guiding aquaculture development sustainably within social-ecological systems^36,37^, and to foster the “ecological well-being”^38^. Yet, few studies have combined CC and EAA principles in their modelling^24–26^ or decision-making processes^39–41^. In the following, we will discuss potentially far-reaching effects of marine aquaculture that are currently underrepresented in the CC debate but crucial to ocean health.

### Moving ahead: embracing spatial ocean connectivity

CC estimates and state indicator assessments have often been based on box system models and applied to small-scale ecosystems well below 500 km^2^ (i.e. estimating CC at bay scale; **Table 1**). This focus on particular settings has its methodological-conceptual reasoning. Further, place-based boundary values are easier to apply in decision-making processes. Nevertheless, the fluidity of space in the ocean allows for passive **dispersal** of organisms and their developmental stages and other aquaculture related output over hundreds of kilometers^42^ (**Figure 1**). As a result of natural **spatial connectivity**, biotic and abiotic entities on which state indicators are based, may ultimately leave areas conventionally considered to estimate CC or even cross international borders (**Figure 1**).

**Table 1:** Examples of state indicators used in different case studies aiming to define physical, production, ecological, assimilative or social aquaculture carrying capacity (CC) for systems of different sizes (following Inglis et al. 2000^16^ in their CC categories, adding *assimilative CC sensu* Chamberlain et al. 2006^28^ and Tett et al. 2011^14^).

| System carrying capacity (CC) | State indicator | Case study reference | Considered area (size) |
| --- | --- | --- | --- |
| Physical CC | Availability of space considering distance from shore and water depth | Yigit et al. 2021<sup>21</sup> | Sigacik Bay, Turkey (0.389 km<sup>2</sup>) |
| Production CC | Depletion of phytoplankton and oxygen | Uribe & Blanco 2001<sup>22</sup> | Tongoy Bay, Chile (55.86 km<sup>2</sup>) |
| Ecological CC | Change of major energy fluxes or structure of the food web | Jiang & Gibbs 2005<sup>23</sup> | Golden and Tasman Bays, New Zealand (4500 km<sup>2</sup>) |
|  | Change in biomass of non-farmed species | Byron et al. 2011<sup>26</sup> | Narragansett Bay, USA (355 km<sup>2</sup>) |
|  | Change in biomass of non-farmed species | Byron et al. 2011<sup>25</sup> | Rhode Island, USA (lagoons from 1.5 to 7.8 km<sup>2</sup>) |
|  | Change in biomass of non-farmed species | Kluger et al. 2016<sup>24</sup> | Sechura Bay, Peru (400 km<sup>2</sup>) |
|  | Depletion of seston | Filgueira et al. 2021<sup>47</sup> | Sober Island, Wine Harbour and Whitehead, Canada (0.9-14.2 km<sup>2</sup>) |
| Assimilative CC | Organic matter, nutrient and contaminant levels | Chamberlain et al. 2006<sup>28</sup> | Great Entry Lagoon, Magdalen Islands, Canada (2.5 km<sup>2</sup>) |
|  | Biodeposition | Weise et al. 2009<sup>29</sup> | Cascapedia Bay and<br>Magdalen Islands,<br>Canada<br>(1.25-2.5 km<sup>2</sup>) |
|  | Waste assimilative<br>capacity and resupply of<br>oxygen | Tett et al. 2011<sup>14</sup> | Loch Creran, Scotland<br>(11 km<sup>2</sup>) |
| Social CC | Satisfaction, acceptability,<br>desirability, preference | Dalton et al. 2017<sup>31</sup> | Narragansett Bay,<br>USA (355 km<sup>2</sup>) |

**Figure 1:**
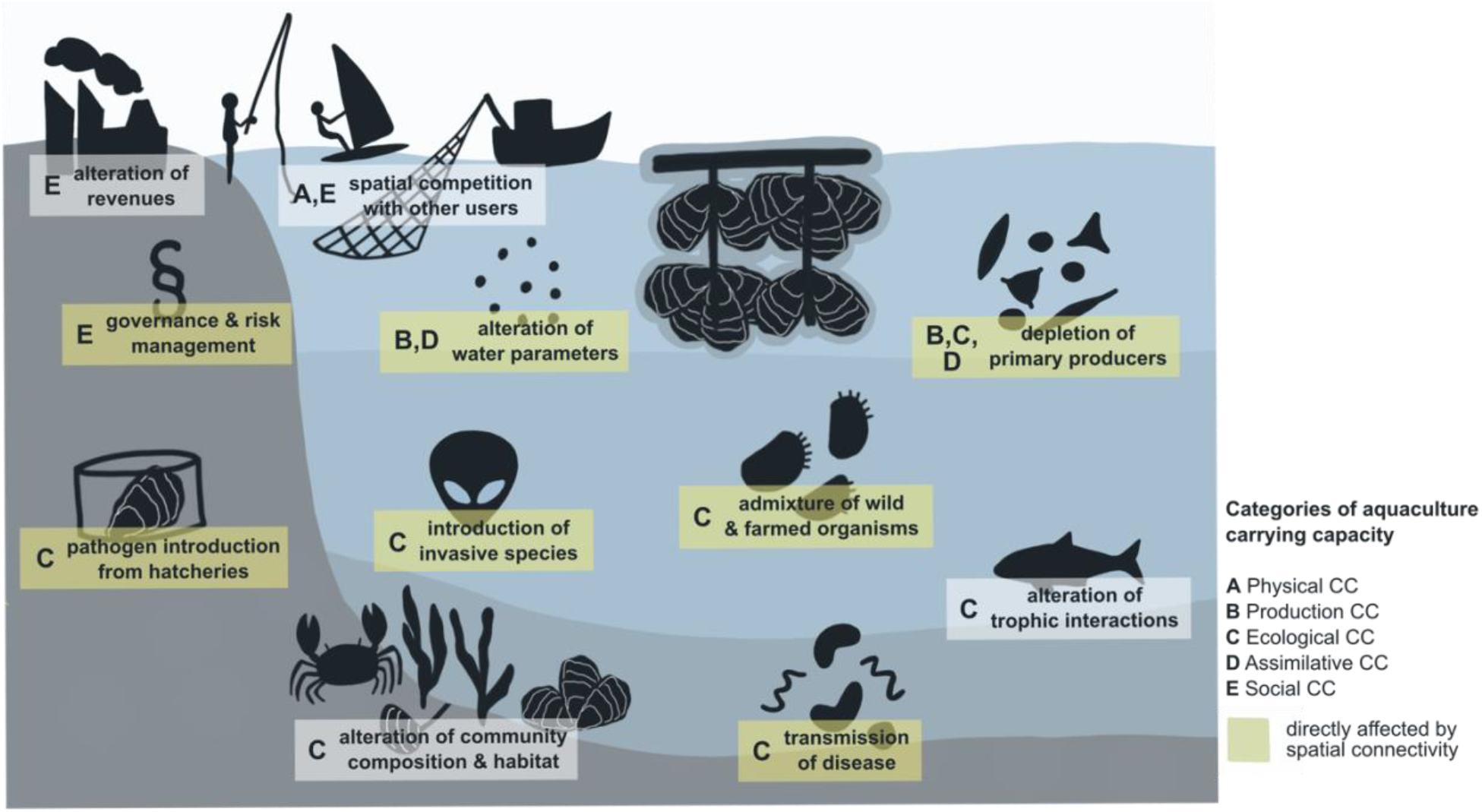
Interactions of marine aquaculture with the surrounding ecosystem and anthropogenic activities that become relevant as state indicators for different aquaculture CC categories (letters). Here the focus is on the example of coastal bivalve aquaculture as one of the first systems to which the concept of aquaculture CC was applied. Almost all state indicators are potentially affected by spatial connectivity within the ocean and hence have effects beyond the typical area considered for CC (highlighted in yellow).

While spatial connectivity in the ocean might seem obvious, the extend of its implications may currently not be fully acknowledged. Also in conservation planning, connectivity has been recently highlighted as a necessary factor^43,44^, importantly on the management and governance level^45^. Few studies so far have considered connectivity in the CC context as the inflow or replenishment of oxygen or seston into the aquaculture area^46,47^. Still, the dispersal away from aquaculture sites is currently largely overlooked. Therefore, the sustainable management of ecosystem services requires a more holistic approach^48^.

### Applying the lens of marine diseases to the carrying capacity debate

In some cases, the dispersal of organisms from aquaculture to the surrounding ecosystem is especially problematic for ocean health and we want to highlight two cases here, which are (i) the dispersal of invasive species and (ii) the dispersal of pathogens or organisms carrying **infectious diseases**. There are several examples of species introduced for economic purposes to a non-native region that have established populations far exceeding their initial point of cultivation with substantial impact on the ecosystem and competition with native species^49^. For example, the Pacific oyster

*Magallana gigas* (previously *Crassostrea gigas*) was introduced to Europe in the 1980s for aquaculture due to their high growth and reproduction rate^50^. Up to now it has established populations all across the North-West European shelf including in places where no oyster aquaculture is pursued^51,52^. Populations continue to expand even against their preferred temperature regime^53^ and most recently into the low saline Baltic Sea^54^. The implications for the environment range from displacement of native species to habitat transformation^55^.

Another highly topical example for the unregulated introduction of non-native species is the seaweed industry^56^ which is at exceptional growth rates in recent years^2,57^. Dispersal of reproductive seaweed material from the initial point of introduction can introduce shifts in the gene pool of spatially connected **wild populations**^10^. Even though numerous cases of non-native species introduction for cultivation are known^58^, translocations and introductions are continuing but urgently need governance and biosecurity regulations to control the translocation of non-native cultivates^56^ as well as the associated diseases^59^.

Together with cultured organisms, also pathogens can escape from aquaculture and disperse to the surrounding environment. Disease outbreaks in aquaculture can greatly affect mortality and growth of farmed animals as well as seafood product quality^30,35^, all of which compromise economic returns. Infection risk is highly dependent on population density^60,61^ with increasing density resulting in an increasing risk of disease transmission. Beyond the economic impact, disease outbreaks and reoccurring mass mortality events are increasingly recognized as a threat to ocean health and expected to intensify in frequency and magnitude under progressing climate change and monoculture practices^62,63^.

Surprisingly, only 24 marine infectious diseases are currently listed by the World Organization for Animal Health (in comparison to 172 terrestrial diseases, March 2023) with major diseases such as the ostreid herpesvirus Os-HV-1 missing. In oyster aquaculture, Os-HV-1 can cause a loss of up to 100% and mass mortalities have already been recorded worldwide^64,65^. However, infections keep recurring due to a limited understanding of the biology of the diseases as well as insufficient management and governance^66^. This fragmentation of both knowledge on and documentation of diseases highlights the challenges imposed to correctly document diseases in the marine realm.

Beyond disease outbreaks within aquaculture farms and the costs involved, there is a high risk for disease spill-over to the surrounding ecosystem (**Figure 1**). Dense populations kept in aquaculture farms can turn into a hub for disease outbreaks transmitted to wild populations^67,68^. The spillover of sea lice from salmon farms, for example, was identified as a reason for wild population declines and local extinction of pink salmon on the Westcoast of Canada^69^. Disease transmission can happen by direct contact to infected organisms or via pathogens shed into seawater where they remain infectious for usually for a short time. Sea lice were shown to cross-infect even distant salmon farms via dispersal by ocean currents^70–72^. Infection of neighboring wild populations is a likely consequence especially in open aquaculture systems, where water exchange between farm and environment is continuous and unavoidable. However, there are also ways to reduce the risk of disease dispersal by considering epidemiology^73^. For example, it was predicted that harvesting oysters prior to the peak disease release season would limit the impact on surrounding organisms^74^. Overall, wild populations are often much less monitored for disease outbreaks than farmed animals due to inaccessibility and hence costs and effort involved, or simply because monitoring of the natural environment is not prioritized by management. Even if infected wild animals are detected, the extend of related mortality events are often unknown and the impact of spill-over from and to aquaculture is not yet fully understood for most systems^75^.

What we do know, is that there are multiple dispersal pathways between farms and the direct and distant environment that are relevant for connectivity and potentially for disease transmission. Farmed bivalves and wild populations are connected via ocean currents (#1 in **Figure 2**) where wild bivalves can potentially act as stepping stones in-between farms (#2) or enhance disease transmission when being in close proximity to farms through spill-over and spill-back of diseases (#3). Human action may have an even bigger impact on spatial connectivity through **anthropogenic dispersal**: transport and relocation of bivalves (and their diseases) (#4,#5), as well as introduction from hatcheries (#6) are vectors for dispersal, even if produced bivalves are consumed locally (#7). Ship transport and ballast water (#8) are additional routes for disease dispersal in the ocean and increase spatial connectivity in the ocean where they can act against direction of ocean currents. Thereby, even very distant locations can in theory be at risk through combined natural and anthropogenic dispersal processes (#9). Considering various dispersal routes and their potential impact on close-by and distant ecosystems, we call to take the resulting multidimensional spatial connectivity into account when providing CC estimates. For example, when applying the lens of marine diseases to CC, options to predict but more importantly reduce the spreading of diseases introduced and amplified by aquaculture should necessarily be developed as to correctly assess the impact of aquaculture on ocean health. Our ocean’s health is, in the end, a combinate product of all ocean’s – and its subsystems – health and the sustainable use of ocean resources inevitably depends thereof.

**Figure 2:**
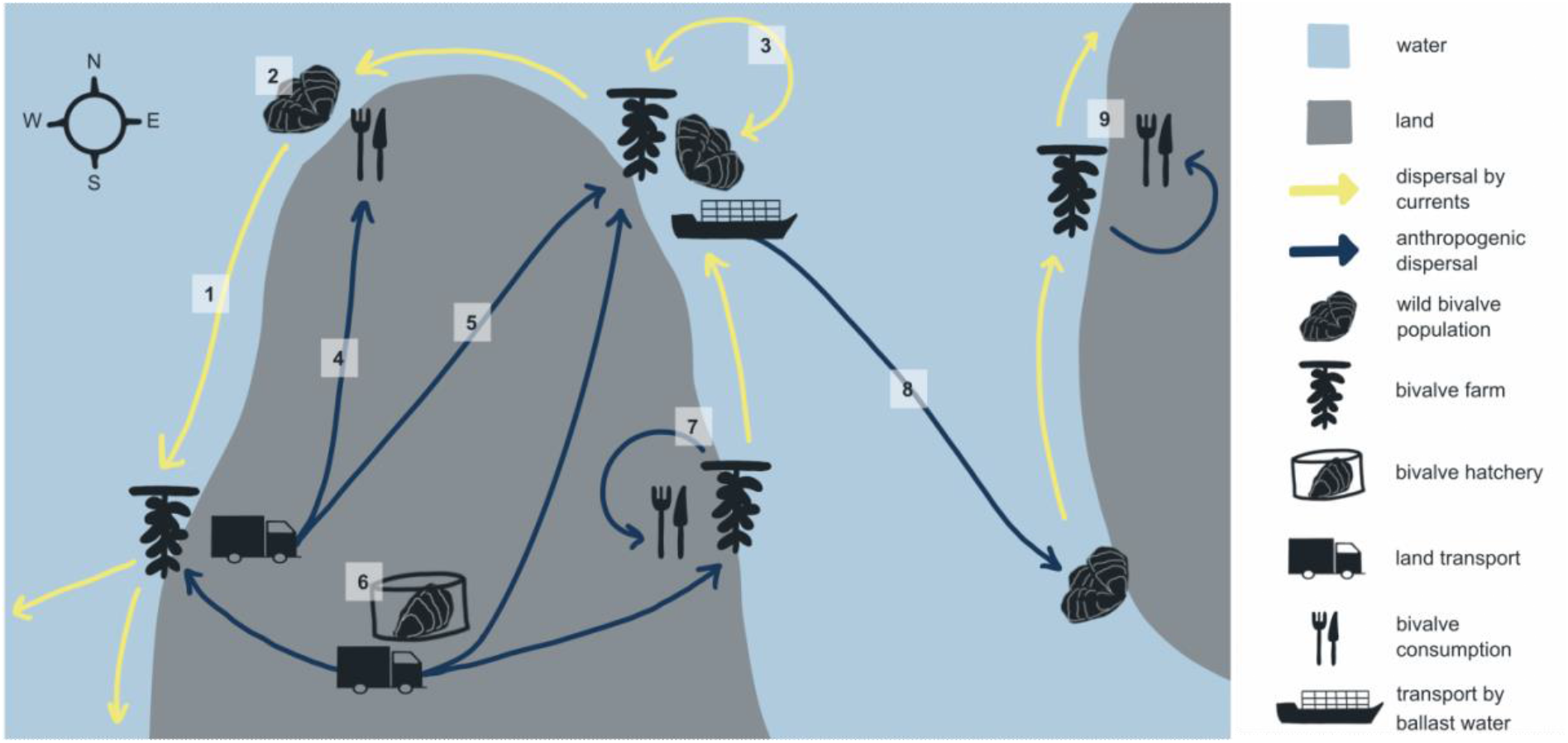
Long-distance dispersal of pathogens as well as farmed organisms introducing connectivity. Dispersal can be passive with ocean currents (in yellow), or via anthropogenic transport (in blue) from hatcheries to different field sites or relocation between sites. Scenarios depicted in the graphic: 1. Connectivity between wild population and farmed bivalves via ocean currents and potential way of disease dispersal. 2. Transport of bivalves (and disease) from production to consumption site. 3. Relocation between different farms. 4. Bivalves (and disease) transported from hatchery to different farms. 5. Local bivalve production and consumption. 6. Wild bivalve population as stepping stone between farms, connected via ocean currents. 7. Wild bivalve population in close proximity to farm. Site of disease spill-over and spill-back. 8. Dispersal of bivalves (and disease) with ship ballast water, also against direction of ocean currents. 9. Bivalves that are only produced and consumed locally are still at risk of infection by the combination of anthropogenic and natural dispersal routes.

### Opportunities for holistic carrying capacity estimates

In the EAA, it is differentiated between several spatial levels to which considerations should apply: farms, watersheds and the global level, emphasizing CC is mostly applied to the farm level^37^. As a first step towards the integration of spatial connectivity in future CC estimates, system boundaries should be carefully rethought. The aim would be to continue providing CC recommendations for the immediately affected and well characterized local area around an aquaculture farm as was done so far, to then predict the potential maximum impact area that is spatially connected.

The strength of connectivity between sites depends on farm size and number of farmed individuals, but also ocean currents and anthropogenic activities such as distance to the next harbor. Continuing with the bivalve aquaculture example (**Figure 1 and 2**), guiding questions for defining system boundaries could be: How many farms are located in that physical space? Where and how large are wild populations of the farmed organisms? Where do seeds come from? Are animals relocated during the growth period? Is there shipping traffic (ballast water exchange) in close proximity? How is the local current regime? Does it change with season? How do ocean currents and their physical-chemical parameters shape connectivity relevant to CC estimates?

Providing quantitative and comprehensive CC estimates is already a complex endeavor at a local scale and valid concerns and constraints have been discussed previously^15,18,35,76,77^. However, especially with respect to governance and international legislation, there should be an interest to consider the potential far-reaching impact of marine aquaculture including those across international boundaries. One potential method to embrace the natural spatial connectivity by ocean currents are hydrodynamic modelling approaches and Lagrangian particle dispersal simulations^78,79^. This toolkit from physical oceanography has been successfully applied to biologically motivated questions, among them the dispersal of passively drifting larvae of various organisms^80,81^ (see **Figure 3 A+B** for more details on the method). Few studies, however, have yet applied dispersal modelling to the spreading of pathogens from aquaculture with ocean currents. One of the few examples is a case study on the infection on sea lice infections in salmon aquaculture farms that demonstrated that increasing the distance between farms would lower the risk of cross-infections^71,72,82^. Although the importance of diseases has been discussed for the context of aquaculture CC earlier^35^, it has only been recently used as a tool to inform CC decisions^83^. In a specific case study, dispersal models were used to assess connectivity and risk of disease dispersal between in- and offshore aquaculture^30^. And while this is a very relevant start to discuss effects of cultures across farm-system-boundaries, the connectivity was pre-defined (in-offshore aquaculture facilities) and other dimensions of spatial connectivity as discussed in the present work were consequently not included.

**Figure 3:**
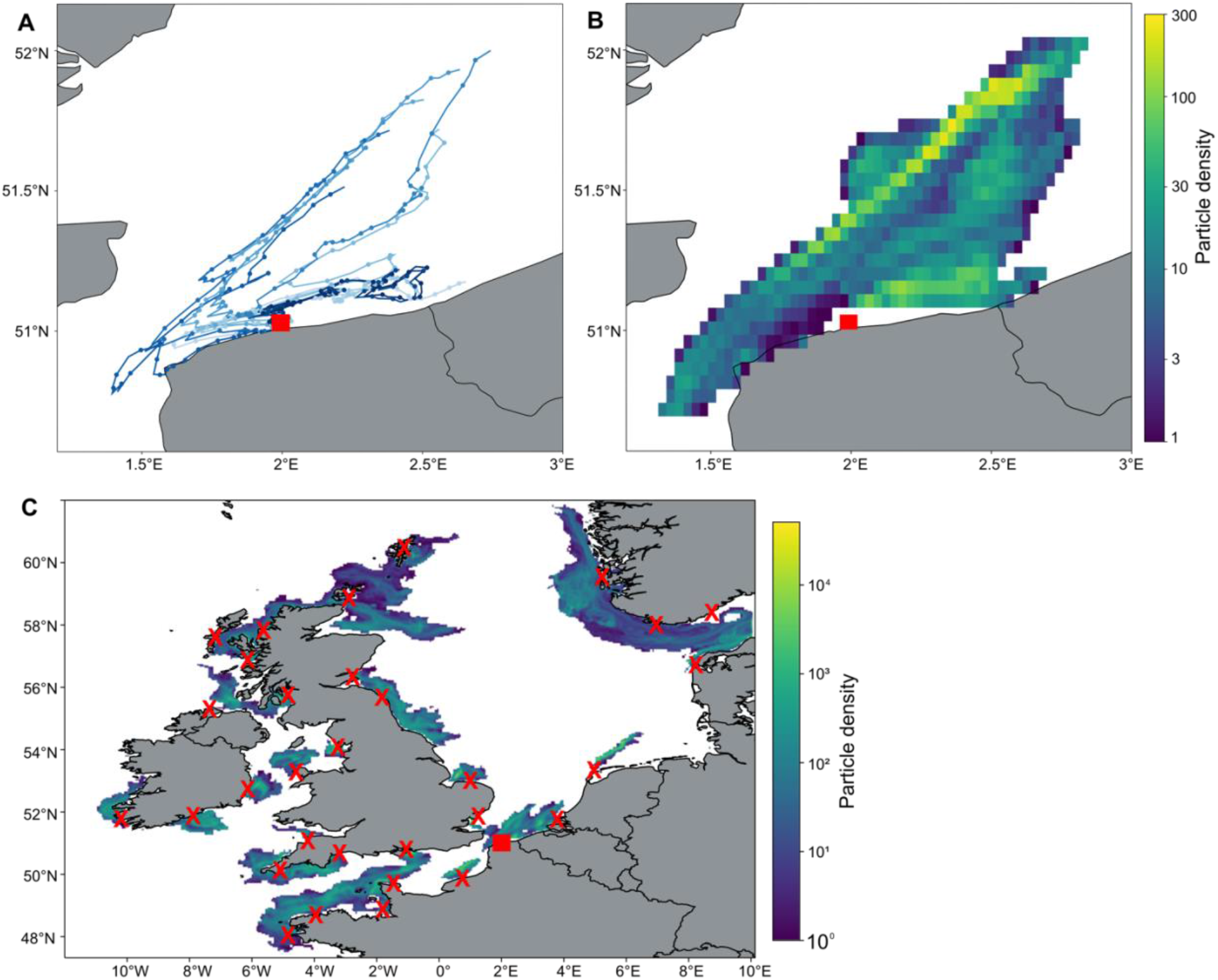
Hydrodynamic ocean models and particle dispersal analysis can provide insights into the potential impact of aquaculture farms through simulation of passive dispersal of farmed organisms and/or associated pathogens. In total, 10,000 virtual particles representing infected oyster larvae were released from an aquaculture farm site on the French North Atlantic coast (red square) during the reproductive season (on 01.07. for the years 2019-2022) in 4 m water depth. The position of virtual particles was tracked every 12 h for 28 days using a km-scale operational ocean model from Copernicus Marine Service^85^. (**A**) Ten representative dispersal trajectories of virtual particles are displayed for the year 2019 (points represent position every 12 h). (**B**) The particle density from day 14-28 after particle release represents the most likely distribution of larvae during the time window for settlement. Larval development time depends on the species but also environmental conditions, and 28 days are within the range of Pacific oyster development^86^. The example shows that ocean currents can disperse infected larvae into different directions along the coast, but also transport them offshore. On average, in 28 days larvae are transported 66.5 km from this example source site in France with a maximum distance of 134.8 km (**Table 2**). Interannual variability of dispersal ranges from 40.1-85.0 km. (**C**) On a larger scale, different locations of farms can be explored as shown here for randomly chosen oyster farms across the North-West European shelf (locations exported from EMODnet^52^ and marked as red X’s. The square marks the example site from panels A and B).

Here, we demonstrate for the example of bivalve aquaculture, the potential impact area of oyster farms at the North West European shelf (**Figure 3 C**, locations of a small subset of farms extracted from EMODnet, accessed January 2023). The simulated entities in this example are infected oyster larvae that pose a risk for disease spread between farms and spill-over to wild populations. On average, infected oyster larvae are predicted to spread over 70 km from aquaculture farms which exceeds most of the areas used to calculate CC (**Table 1**). Depending on the exact location of farms, their proximity to the coast and exposure to the open ocean, dispersal distances can vary greatly. Even in this rather small dataset, average dispersal varies between 21 and over 200 km, demonstrating the need and benefit of applying dispersal simulations to each specific case. An advantage over, for example, genetic methods of assessing spatial connectivity between populations, is the potential for high-throughput and fast application to several locations at a time. Further, simulations are non-invasive and can be applied without much *a priori* knowledge of a region if a hydrodynamic model is available (e.g., Copernicus Marine Service provides an open access platform for various hydrodynamic models of the global ocean^84^).

**Table 2:**
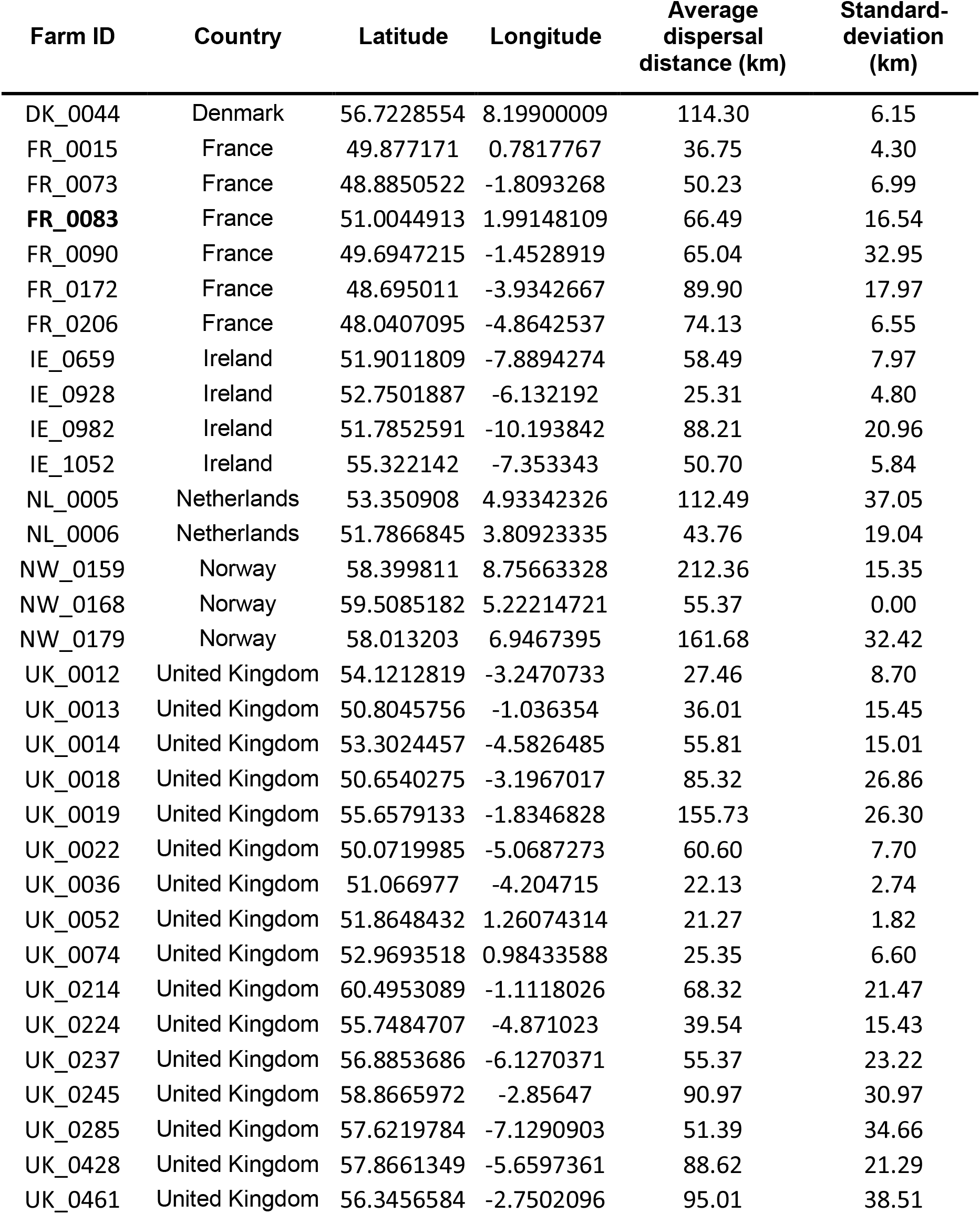
Average dispersal distance of simulated oyster larvae over 28 days in July 2019-2022 from aquaculture farms across the North-West European shelf (**Figure 3**). Example station from Figure 3 A+B is highlighted in bold.

| Farm ID | Country | Latitude | Longitude | Average dispersal distance (km) | Standard-deviation (km) |
| --- | --- | --- | --- | --- | --- |
| DK_0044 | Denmark | 56.7228554 | 8.19900009 | 114.30 | 6.15 |
| FR_0015 | France | 49.877171 | 0.7817767 | 36.75 | 4.30 |
| FR_0073 | France | 48.8850522 | -1.8093268 | 50.23 | 6.99 |
| **FR_0083** | France | 51.0044913 | 1.99148109 | 66.49 | 16.54 |
| FR_0090 | France | 49.6947215 | -1.4528919 | 65.04 | 32.95 |
| FR_0172 | France | 48.695011 | -3.9342667 | 89.90 | 17.97 |
| FR_0206 | France | 48.0407095 | -4.8642537 | 74.13 | 6.55 |
| IE_0659 | Ireland | 51.9011809 | -7.8894274 | 58.49 | 7.97 |
| IE_0928 | Ireland | 52.7501887 | -6.132192 | 25.31 | 4.80 |
| IE_0982 | Ireland | 51.7852591 | -10.193842 | 88.21 | 20.96 |
| IE_1052 | Ireland | 55.322142 | -7.353343 | 50.70 | 5.84 |
| NL_0005 | Netherlands | 53.350908 | 4.93342326 | 112.49 | 37.05 |
| NL_0006 | Netherlands | 51.7866845 | 3.80923335 | 43.76 | 19.04 |
| NW_0159 | Norway | 58.399811 | 8.75663328 | 212.36 | 15.35 |
| NW_0168 | Norway | 59.5085182 | 5.22214721 | 55.37 | 0.00 |
| NW_0179 | Norway | 58.013203 | 6.9467395 | 161.68 | 32.42 |
| UK_0012 | United Kingdom | 54.1212819 | -3.2470733 | 27.46 | 8.70 |
| UK_0013 | United Kingdom | 50.8045756 | -1.036354 | 36.01 | 15.45 |
| UK_0014 | United Kingdom | 53.3024457 | -4.5826485 | 55.81 | 15.01 |
| UK_0018 | United Kingdom | 50.6540275 | -3.1967017 | 85.32 | 26.86 |
| UK_0019 | United Kingdom | 55.6579133 | -1.8346828 | 155.73 | 26.30 |
| UK_0022 | United Kingdom | 50.0719985 | -5.0687273 | 60.60 | 7.70 |
| UK_0036 | United Kingdom | 51.066977 | -4.204715 | 22.13 | 2.74 |
| UK_0052 | United Kingdom | 51.8648432 | 1.26074314 | 21.27 | 1.82 |
| UK_0074 | United Kingdom | 52.9693518 | 0.98433588 | 25.35 | 6.60 |
| UK_0214 | United Kingdom | 60.4953089 | -1.1118026 | 68.32 | 21.47 |
| UK_0224 | United Kingdom | 55.7484707 | -4.871023 | 39.54 | 15.43 |
| UK_0237 | United Kingdom | 56.8853686 | -6.1270371 | 55.37 | 23.22 |
| UK_0245 | United Kingdom | 58.8665972 | -2.85647 | 90.97 | 30.97 |
| UK_0285 | United Kingdom | 57.6219784 | -7.1290903 | 51.39 | 34.66 |
| UK_0428 | United Kingdom | 57.8661349 | -5.6597361 | 88.62 | 21.29 |
| UK_0461 | United Kingdom | 56.3456584 | -2.7502096 | 95.01 | 38.51 |

In order to integrate this method into CC assessments, thresholds of acceptable connectivity would need to be assessed and defined. This might be achieved by setting a limit to a “maximum impact area” or “maximum impact distance” depending on the epidemiology of a disease. Beyond the use for site selection for aquaculture, connectivity could be re-assessed regularly, since it depends not only on local current conditions but also on weather and other environmental variables and can vary between years. As such, simulations might be used to determine annual production limits or harvesting times. Even in the event of a disease outbreak within a farm, simulations could be used to inform potentially affected farms or protected areas to increase monitoring efforts. Especially farms close to international borders or other legislative districts are expected to benefit from early warning systems (**Figure 3 B**).

We can only predict, not change ocean currents and thus have to embrace their flow and the spatial connectivity they introduce into our considerations. However, we can certainly influence anthropogenic dispersal of farmed organisms and their pathogens and reducing the latter should always be a priority. In ecological terms: the less anthropogenic transport the better, though not always avoidable. Further, a well-documented disease screening of both native and non-native organisms before introduction to aquaculture sites is absolutely crucial to reduce pathogen loading and spill-over of diseases. Similarly, a regular disease screening of already cultivated organisms ensures early detection before mortality events occur and thus stable biomass production and profit. Optimizing the timing of harvest of farmed oysters as to reduce the pathogen loading of the environment may be a management option, though the costs for producers has to be priced in. Lastly, any recorded and quantified disease breakouts should be publicly available, best in an international database, providing a resource for early detection and warning system before disease spread occurs. Overall, the design and implementation of meaningful regulations is the most important measure to control unwanted introductions of non-native species and their potential pathogens for aquaculture purposes.

## Summary

1. Aquaculture CC calculations need to embrace spatial connectivity in the ocean and the potential far reaching effects through dispersal by ocean currents.
2. Embracing spatial connectivity in CC calculations allows to improve estimations of aquaculture impact on the environment and ocean health that currently might be underestimated.
3. Spatial connectivity touches several impacts of aquaculture on ecosystems, while the introduction and dispersal of invasive species and marine diseases are of highest concern.
4. Hydrodynamic modelling and Lagrangian particle dispersal simulations provide tools to assess the potential area affected by marine aquaculture and predict connectivity to distant ecosystems and other farms including those across international boundaries.
5. Importantly, simulations are suggested as part of a multicriteria approach and not to replace current ecological or social CC frameworks.
6. Understanding dispersal from aquaculture farms is a first step towards limiting it actively to a minimum.
7. Including connectivity and the focus on ocean health into the CC concept has implications for natural, social, political and economic sciences as well as governance. It provides a crucial step towards sustainably promoting aquaculture as to feed the ever-growing human population, while maintaining and protecting ocean health.

## Acknowledgements

We would like to thank the organizers of the 2^nd^ Ocean Health Symposium Kiel for creating an inspiring event fostering discussions and formation of interdisciplinary collaborations. We are grateful to Arne Biastoch, Sophie Koch and Tyler Carrier for their feedback on an earlier version of this manuscript. Thanks to the students Jana Noller and Leon Mock for their support on figures and tables. We also appreciate the friendly contact to the editorial team of *One Earth*. This study has been conducted using E.U. Copernicus Marine Service Information; https://doi.org/10.48670/moi-00054

## References

1. Franke, A., Blenckner, T., Duarte, C.M., Ott, K., Fleming, L.E., Antia, A., Reusch, T.B.H., Bertram, C., Hein, J., Kronfeld-Goharani, U., et al. (2020). Operationalizing Ocean Health: Toward Integrated Research on Ocean Health and Recovery to Achieve Ocean Sustainability. One Earth 2, 557–565. 10.1016/j.oneear.2020.05.013.

2. FAO: The State of World Fisheries and Aquaculture 2022. Towards Blue Transformation (2022). 10.4060/cc0461en.

3. Edwards, P. (2015). Aquaculture environment interactions: Past, present and likely future trends. Aquaculture 447, 2–14. 10.1016/j.aquaculture.2015.02.001.

4. Ottinger, M., Clauss, K., and Kuenzer, C. (2016). Aquaculture: Relevance, distribution, impacts and spatial assessments – A review. Ocean. Coast. Manag. 119, 244–266. 10.1016/j.ocecoaman.2015.10.015.

5. Gallardi, D. (2014). Effects of Bivalve Aquaculture on the Environment and Their Possible Mitigation: A Review. Fish. Aquac. J. 5, 1000105. 10.4172/2150-3508.1000105.

6. Hulot, V., Saulnier, D., Lafabrie, C., and Gaertner-Mazouni, N. (2020). Shellfish culture: a complex driver of planktonic communities. Rev. Aquacult. 12, 33–46. 10.1111/raq.12303.

7. Liu, X., and Zhang, X. (2022). Impacts of High-Density Suspended Aquaculture on Water Currents: Observation and Modeling. JMSE 10, 1151. 10.3390/jmse10081151.

8. Liu, Z., Ma, A., Mathé, E., Merling, M., Ma, Q., and Liu, B. (2021). Network analyses in microbiome based on high-throughput multi-omics data. Briefings in bioinformatics 22, 1639–1655. 10.1093/bib/bbaa005.

9. Ye, L.-X., Ritz, D.A., Fenton, G.E., and Lewis, M.E. (1991). Tracing the influence on sediments of organic waste from a salmonid farm using stable isotope analysis. J. Exp. Mar. Biol 145, 161–174. 10.1016/0022-0981(91)90173-T.

10. Tano, S.A., Halling, C., Lind, E., Buriyo, A., and Wikström, S.A. (2015). Extensive spread of farmed seaweeds causes a shift from native to non-native haplotypes in natural seaweed beds. Mar. Biol. 162, 1983–1992. 10.1007/s00227-015-2724-7.

11. Mckindsey, C.W., Landry, T., O’Beirn, F.X., and Davies, I.M. (2007). Bivalve aquaculture and exotic species: A review of ecological considerations and management issues. J. Shellfish Res. 26, 281–294. 10.2983/0730-8000(2007)26[281:BAAESA]2.0.CO;2.

12. Van Wesenbeeck, B.K., Balke, T., Van Eijk, P., Tonneijck, F., Siry, H.Y., Rudianto, M.E., and Winterwerp, J.C. (2015). Aquaculture induced erosion of tropical coastlines throws coastal communities back into poverty. Ocean. Coast. Manag. 116, 466–469. 10.1016/j.ocecoaman.2015.09.004.

13. Halpern, B.S., Longo, C., Hardy, D., McLeod, K.L., Samhouri, J.F., Katona, S.K., Kleisner, K., Lester, S.E., O’Leary, J., Ranelletti, M., et al. (2012). An index to assess the health and benefits of the global ocean. Nature 488, 615–620. 10.1038/nature11397.

14. Tett, P., Portilla, E., Gillibrand, P.A., and Inall, M. (2011). Carrying and assimilative capacities: the ACExR-LESV model for sea-loch aquaculture: Carrying and assimilative capacities. Aquac. Res. 42, 51–67. 10.1111/j.1365-2109.2010.02729.x.

15. McKindsey, C.W., Thetmeyer, H., Landry, T., and Silvert, W. (2006). Review of recent carrying capacity models for bivalve culture and recommendations for research and management. Aquaculture 261, 451–462. 10.1016/j.aquaculture.2006.06.044.

16. Inglis, G.J., Hayden, B.J., and Ross, A.H. (2000). An overview of factors affecting the carrying capacity of coastal embayments for mussel culture. NIWA Client Report CHC00/69, Christchurch, New Zealand.

17. Chapman, E.J., and Byron, C.J. (2018). The flexible application of carrying capacity in ecology. Glob. Ecol. Conserv 13, e00365. 10.1016/j.gecco.2017.e00365.

18. Weitzman, J., and Filgueira, R. (2019). The evolution and application of carrying capacity in aquaculture: towards a research agenda. Rev. Aquacult 12, 1297– 1322. 10.1111/raq.12383.

19. Ross, L.G., Telfer, T., Falconer, L., Soto, D., and Aguilar-Manjarrez, J. eds. (2013). Site selection and carrying capacities for inland and coastal aquaculture (Food and Agriculture Organization of the United Nations).

20. Teixeira, Z., Marques, C., Mota, J.S., and Garcia, A.C. (2018). Identification of potential aquaculture sites in solar saltscapes via the Analytic Hierarchy Process. Ecol. Indic 93, 231–242. 10.1016/j.ecolind.2018.05.003.

21. Yigit, M., Ergun, S., Buyukates, Y., Ates, A.S., and Ozdilek, H.G. (2021). Physical carrying capacity of a potential aquaculture site in the Mediterranean: the case of Sigacik Bay, Turkey. Environ. Sci. Pollut. Res 28, 9753–9759. 10.1007/s11356-020-11455-y.

22. Uribe, E., and Blanco, Y.J.L. (2001). 12: Carrying capacity of aquatic systems to support intensive scallop aquaculture; the case of Argopecten purpuratus, Bahia Tongoy, Chile. In Los Moluscos Pectínidos de Iberoamérica: Ciencia y Acuicultura (Maeda-Martinez, A.N.).

23. Jiang, W., and Gibbs, M.T. (2005). Predicting the carrying capacity of bivalve shellfish culture using a steady, linear food web model. Aquaculture 244, 171–185. 10.1016/j.aquaculture.2004.11.050.

24. Kluger, L.C., Taylor, M.H., Mendo, J., Tam, J., and Wolff, M. (2016). Carrying capacity simulations as a tool for ecosystem-based management of a scallop aquaculture system. Ecol. Model 331, 44–55. 10.1016/j.ecolmodel.2015.09.002.

25. Byron, C., Link, J., Costa-Pierce, B., and Bengtson, D. (2011). Modeling ecological carrying capacity of shellfish aquaculture in highly flushed temperate lagoons. Aquaculture 314, 87–99. 10.1016/j.aquaculture.2011.02.019.

26. Byron, C., Link, J., Costa-Pierce, B., and Bengtson, D. (2011). Calculating ecological carrying capacity of shellfish aquaculture using mass-balance modeling: Narragansett Bay, Rhode Island. Ecol. Model 222, 1743–1755. 10.1016/j.ecolmodel.2011.03.010.

27. Kluger, L.C., Filgueira, R., and Wolff, M. (2017). Integrating the Concept of Resilience into an Ecosystem Approach to Bivalve Aquaculture Management. Ecosystems 20, 1364–1382. 10.1007/s10021-017-0118-z.

28. Chamberlain, J., Weise, A.M., Dowd, M., and Grant, J. (2006). Modeling approaches to assess the potential effects of shellfish aquaculture on the marine environment. Fisheries and Oceans Canada.

29. Weise, A.M., Cromey, C.J., Callier, M.D., Archambault, P., Chamberlain, J., and McKindsey, C.W. (2009). Shellfish-DEPOMOD: Modelling the biodeposition from suspended shellfish aquaculture and assessing benthic effects. Aquaculture 288, 239–253. 10.1016/j.aquaculture.2008.12.001.

30. Ferreira, J.G., Saurel, C., Lencart e Silva, J.D., Nunes, J.P., and Vazquez, F. (2014). Modelling of interactions between inshore and offshore aquaculture. Aquaculture 426–427, 154–164. 10.1016/j.aquaculture.2014.01.030.

31. Dalton, T., Jin, D., Thompson, R., and Katzanek, A. (2017). Using normative evaluations to plan for and manage shellfish aquaculture development in Rhode Island coastal waters. Mar. Policy 83, 194–203. 10.1016/j.marpol.2017.06.010.

32. Byron, C.J., Jin, D., and Dalton, T.M. (2015). An Integrated ecological–economic modeling framework for the sustainable management of oyster farming. Aquaculture 447, 15–22. 10.1016/j.aquaculture.2014.08.030.

33. Kluger, L.C., and Filgueira, R. (2021). Thinking outside the box: embracing social complexity in aquaculture carrying capacity estimations. ICES J. Mar. Sci 78, 435– 442. 10.1093/icesjms/fsaa063.

34. Byron, C.J., and Costa-Pierce, B.A. (2013). Carrying capacity tools for use in the implementation of an ecosystems approach to aquaculture. In Site selection and carrying capacities for inland and coastal aquaculture FAO Fisheries and Aquaculture Proceedings., pp. 87–101.

35. Ferreira, J.G., Ramos, L., and Costa-Pierce, B.A. (2013). Key drivers and issues surrounding carrying capacity and site selection, with emphasis on environmental components. In Site selection and carrying capacities for inland and coastal aquaculture FAO Fisheries and Aquaculture Proceedings., pp. 47–86.

36. Brugère, C., Aguilar-Manjarrez, J., Beveridge, M.C.M., and Soto, D. (2019). The ecosystem approach to aquaculture 10 years on - a critical review and consideration of its future role in blue growth. Rev. Aquacult 11, 493–514. 10.1111/raq.12242.

37. Soto, D., Aguilar-Manjarrez, J., Brugère, C., Angel, Bailey, C., Black, K., Edwards, P., Costa-Pierce, B., Chopin, T., Deudero, S., et al. (2008). Applying an ecosystem-based approach to aquaculture: principles, scales and some management measures. In FAO Fisheries and Aquaculture Proceedings, Soto, D., Aguilar-Manjarrez, J., Hishamunda, N., ed., pp. 15–35.

38. FAO (2020). Ecosystem Approach to Aquaculture Management - Handbook. https://www.fao.org/policy-support/tools-and-publications/resources-details/en/c/1401156/.

39. Silva, C., Barbieri, M.A., Yanez, E., Gutierrez, J.C., and Angel DelValls, E.T. (2012). Using indicators and models for an ecosystem approach to fisheries and aquaculture management: the anchovy fishery and Pacific oyster culture in Chile: case studies. lajar 40, 955–969. 10.3856/vol40-issue4-fulltext-12.

40. Hermawan, S. (2018). The Benefit of Decision Support System as Sustainable Environment Technology to Utilize Coastal Abundant Resources in Indonesia. MATEC Web Conf. 164, 01043. 10.1051/matecconf/201816401043.

41. Fisher, J., Angel, D., Callier, M., Cheney, D., Filgueira, R., Hudson, B., McKindsey, C.W., Milke, L., Moore, H., O’Beirn, F., et al. (2023). Ecological carrying capacity in mariculture: Consideration and application in geographic strategies and policy. Mar. Policy 150, 105516. 10.1016/j.marpol.2023.105516.

42. Kinlan, B.P., and Gaines, S.D. (2003). Propagule dispersal in marine and terrestrial environments: a community perspective. Ecology 84, 2007–2020. 10.1890/01-0622.

43. Beger, M., Metaxas, A., Balbar, A.C., McGowan, J.A., Daigle, R., Kuempel, C.D., Treml, E.A., and Possingham, H.P. (2022). Demystifying ecological connectivity for actionable spatial conservation planning. Trends Ecol. Evol 37, 1079–1091. 10.1016/j.tree.2022.09.002.

44. Legrand, T., Chenuil, A., Ser-Giacomi, E., Arnaud-Haond, S., Bierne, N., and Rossi, V. (2022). Spatial coalescent connectivity through multi-generation dispersal modelling predicts gene flow across marine phyla. Nat. Commun 13, 5861. 10.1038/s41467-022-33499-z.

45. Keeley, A.T.H., Fremier, A.K., Goertler, P.A.L., Huber, P.R., Sturrock, A.M., Bashevkin, S.M., Barbaree, B.A., Grenier, J.L., Dilts, T.E., Gogol-Prokurat, M., et al. (2022). Governing Ecological Connectivity in Cross-Scale Dependent Systems. BioScience 72, 372–386. 10.1093/biosci/biab140.

46. Filgueira, R., Guyondet, T., Comeau, L.A., and Grant, J. (2014). A fully-spatial ecosystem-DEB model of oyster (Crassostrea virginica) carrying capacity in the Richibucto Estuary, Eastern Canada. J. Mar. Syst 136, 42–54. 10.1016/j.jmarsys.2014.03.015.

47. Filgueira, R., Guyondet, T., Thupaki, P., Sakamaki, T., and Grant, J. (2021). The effect of embayment complexity on ecological carrying capacity estimations in bivalve aquaculture sites. J. Clean. Prod 288, 125739. 10.1016/j.jclepro.2020.125739.

48. Rector, M.E., Filgueira, R., and Grant, J. (2021). Ecosystem services in salmon aquaculture sustainability schemes. Ecosyst. Serv 52. https://doi.org/10.1016/j.ecoser.2021.101379.

49. Castellanos-Galindo, G.A., Moreno, X., and Robertson, R. (2018). Risks to eastern Pacific marine ecosystems from sea-cage mariculture of alien Cobia. MBI 9, 323– 327. 10.3391/mbi.2018.9.3.14.

50. Lallias, D., Boudry, P., Batista, F.M., Beaumont, A., King, J.W., Turner, J.R., and Lapègue, S. (2015). Invasion genetics of the Pacific oyster Crassostrea gigas in the British Isles inferred from microsatellite and mitochondrial markers. Biol. Invasions 17, 2581–2595 10.1007/s10530-015-0896-1.

51. OBIS: Ocean Biodiversity Information System. Magallana gigas https://www.obis.org/taxon/836033.

52. EMODnet Human Activities: Shellfish aquaculture https://ows.emodnet-humanactivities.eu/geonetwork/srv/api/records/aa0d2b45-49c4-4b42-86bb-8971a3c2d2cc.

53. Anglès d’Auriac, M.B., Rinde, E., Norling, P., Lapègue, S., Staalstrøm, A., Hjermann, D.ø., and Thaulow, J. (2017). Rapid expansion of the invasive oyster Crassostrea gigas at its northern distribution limit in Europe: Naturally dispersed or introduced? PLoS ONE 12, e0177481. 10.1371/journal.pone.0177481.

54. Ewers-Saucedo, C., Heuer, N., Moesges, Z., Ovenbeck, K., Schröter, N., and Brandis, D. (2020). First record of the Pacific oyster Magallana gigas (Thunberg, 1793) in the Baltic Sea proper. Mar. Biodivers. Rec 13, 9 10.1186/s41200-020-00193-2.

55. Herbert, R.J.H., Humphreys, J., Davies, Clare.J., Roberts, C., Fletcher, S., and Crowe, Tasman.P. (2016). Ecological impacts of non-native Pacific oysters (Crassostrea gigas) and management measures for protected areas in Europe. Biodivers. Conserv 25, 2835–2865. 10.1007/s10531-016-1209-4.

56. Brakel, J., Sibonga, R.C., Dumilag, R.V., Montalescot, V., Campbell, I., Cottier-Cook, E.J., Ward, G., Le Masson, V., Liu, T., Msuya, F.E., et al. (2021). Exploring, harnessing and conserving marine genetic resources towards a sustainable seaweed aquaculture. Plants People Planet 3, 337–349. 10.1002/ppp3.10190.

57. FAO: The state of the worlds fisheries and aquaculture 2020. Sustainability in action (2020). https://www.fao.org/documents/card/en/c/ca9229en.

58. Molnar, J.L., Gamboa, R.L., Revenga, C., and Spalding, M.D. (2008). Assessing the global threat of invasive species to marine biodiversity. Front. Ecol. Environ 6, 485–492. 10.1890/070064.

59. Campbell, I., Kambey, C.S.B., Mateo, J.P., Rusekwa, S.B., Hurtado, A.Q., Msuya, F.E., Stentiford, G.D., and Cottier-Cook, E.J. (2020). Biosecurity policy and legislation for the global seaweed aquaculture industry. J. Appl. Phycol 32, 2133– 2146. 10.1007/s10811-019-02010-5.

60. Jansen, P.A., Kristoffersen, A.B., Viljugrein, H., Jimenez, D., Aldrin, M., and Stien, A. (2012). Sea lice as a density-dependent constraint to salmonid farming. Proc. R. Soc. B 279, 2330–2338. 10.1098/rspb.2012.0084.

61. Krkosek, M. (2010). Host density thresholds and disease control for fisheries and aquaculture. Aquacult. Environ. Interact 1, 21–32. 10.3354/aei0004.

62. Tracy, A.M., Pielmeier, M.L., Yoshioka, R.M., Heron, S.F., and Harvell, C.D. (2019). Increases and decreases in marine disease reports in an era of global change. Proc. R. Soc. B 286, 20191718. 10.1098/rspb.2019.1718.

63. Groner, M.L., Maynard, J., Breyta, R., Carnegie, R.B., Dobson, A., Friedman, C.S., Froelich, B., Garren, M., Gulland, F.M.D., Heron, S.F., et al. (2016). Managing marine disease emergencies in an era of rapid change. Philos. Trans. R. Soc. Lon. B. Biol. Sci 371. 10.1098/rstb.2015.0364.

64. Alfaro, A.C., Nguyen, T.V., and Merien, F. (2019). The complex interactions of Ostreid herpesvirus 1, Vibrio bacteria, environment and host factors in mass mortality outbreaks of Crassostrea gigas. Rev. Aquacult 11, 1148–1168. 10.1111/raq.12284.

65. Burge, C., Griffin, F., and Friedman, C. (2006). Mortality and herpesvirus infections of the Pacific oyster Crassostrea gigas in Tomales Bay, California, USA. Dis. Aquat. Org 72, 31–43. 10.3354/dao072031.

66. Pernet, F., Lupo, C., Bacher, C., and Whittington, R.J. (2016). Infectious diseases in oyster aquaculture require a new integrated approach. Phil. Trans. R. Soc. B 371, 20150213. 10.1098/rstb.2015.0213.

67. Krkošek, M., Lewis, M.A., and Volpe, J.P. (2005). Transmission dynamics of parasitic sea lice from farm to wild salmon. Proc. R. Soc. B 272, 689–696. 10.1098/rspb.2004.3027.

68. Lafferty, K.D., and Ben-Horin, T. (2013). Abalone farm discharges the withering syndrome pathogen into the wild. Front. Microbiol 4. 10.3389/fmicb.2013.00373.

69. Krkošek, M., Ford, J.S., Morton, A., Lele, S., Myers, R.A., and Lewis, M.A. (2007). Declining Wild Salmon Populations in Relation to Parasites from Farm Salmon. Science 318, 1772–1775. 10.1126/science.1148744.

70. Cantrell, D., Vanderstichel, R., Filgueira, R., Grant, J., and Revie, C. (2021). Validation of a sea lice dispersal model: principles from ecological agent-based models applied to aquatic epidemiology. Aquacult. Environ. Interact 13, 65–79. 10.3354/aei00390.

71. Salama, N.K.G., Collins, C.M., Fraser, J.G., Dunn, J., Pert, C.C., Murray, A.G., and Rabe, B. (2013). Development and assessment of a biophysical dispersal model for sea lice. J. Fish Dis 36, 323–337. 10.1111/jfd.12065.

72. Salama, N.K.G., Murray, A.G., and Rabe, B. (2016). Simulated environmental transport distances of Lepeophtheirus salmonis in Loch Linnhe, Scotland, for informing aquaculture area management structures. J. Fish Dis 39, 419–428. 10.1111/jfd.12375.

73. Arzul, I., Furones, D., Cheslett, D., Gennari, L., Delangle, E., Enez, F., Lupo, C., Mortensen, S., Pernet, F., and Peeler, E. (2021). Manual for bivalve disease management and biosecurity - H2020 VIVALDI Project.

74. Ben-Horin, T., Burge, C., Bushek, D., Groner, M., Proestou, D., Huey, L., Bidegain, G., and Carnegie, R. (2018). Intensive oyster aquaculture can reduce disease impacts on sympatric wild oysters. Aquacult. Environ. Interact 10, 557–567. 10.3354/aei00290.

75. Bouwmeester, M.M., Goedknegt, M.A., Poulin, R., and Thieltges, D.W. (2021). Collateral diseases: Aquaculture impacts on wildlife infections. J. Appl. Ecol 58, 453–464. 10.1111/1365-2664.13775.

76. McKindsey, C.W. (2013). Carrying Capacity for Sustainable Bivalve Aquaculture. In Sustainable Food Production, P. Christou, R. Savin, B. A. Costa-Pierce, I. Misztal, and C. B. A. Whitelaw, eds. (Springer New York), pp. 449–466. 10.1007/978-1-4614-5797-8_179.

77. Filgueira, R., Comeau, L.A., Guyondet, T., McKindsey, C.W., and Byron, C.J. (2015). Modelling Carrying Capacity of Bivalve Aquaculture: A Review of Definitions and Methods. In Encyclopedia of Sustainability Science and Technology, R. A. Meyers, ed. (Springer New York), pp. 1–33. 10.1007/978-1-4939-2493-6_945-1.

78. van Sebille, E., Griffies, S.M., Abernathey, R., Adams, T.P., Berloff, P., Biastoch, A., Blanke, B., Chassignet, E.P., Cheng, Y., Cotter, C.J., et al. (2018). Lagrangian ocean analysis: Fundamentals and practices. Ocean Model 121, 49–75. 10.1016/j.ocemod.2017.11.008.

79. Delandmeter, P., and Van Sebille, E. (2019). The Parcels v2.0 Lagrangian framework: new field interpolation schemes. Geosci. Model Dev 12, 3571–3584. 10.5194/gmd-12-3571-2019.

80. Gary, S.F., Fox, A.D., Biastoch, A., Roberts, J.M., and Cunningham, S.A. (2020). Larval behaviour, dispersal and population connectivity in the deep sea. Sci. Rep 10, 1–12. 10.1038/s41598-020-67503-7.

81. Busch, K., Taboada, S., Riesgo, A., Koutsouveli, V., Ríos, P., Cristobo, J., Franke, A., Getzlaff, K., Schmidt, C., Biastoch, A., et al. (2021). Population connectivity of fan-shaped sponge holobionts in the deep Cantabrian Sea. Deep Sea Res. Part I Oceanogr 167, 103427. 10.1016/j.dsr.2020.103427.

82. Cantrell, D.L., Groner, M.L., Ben-Horin, T., Grant, J., and Revie, C.W. (2020). Modeling Pathogen Dispersal in Marine Fish and Shellfish. Trends Parasitol 36, 239–249. 10.1016/j.pt.2019.12.013.

83. Ferreira, J.G., Taylor, N.G.H., Cubillo, A., Lencart-Silva, J., Pastres, R., Bergh, ø., and Guilder, J. (2021). An integrated model for aquaculture production, pathogen interaction, and environmental effects. Aquaculture 536, 736438. 10.1016/j.aquaculture.2021.736438.

84. Copernicus Marine Service https://marine.copernicus.eu/.

85. Copernicus Marine Service: Atlantic - European North West Shelf - Ocean Physics Analysis and Forecast https://doi.org/10.48670/moi-00054.

86. Bayne, B.L. (2017). Reproduction. In Developments in Aquaculture and Fisheries Science (Elsevier), pp. 565–701. 10.1016/B978-0-12-803472-9.00009-1.

